# Strain and rupture of HIV-1 capsids during uncoating

**DOI:** 10.1101/2021.09.30.462583

**Authors:** Alvin Yu, Elizabeth M.Y. Lee, John A.G. Briggs, Barbie K. Ganser-Pornillos, Owen Pornillos, Gregory A. Voth

## Abstract

Viral replication in HIV-1 relies on a fullerene-shaped capsid to transport genetic material deep into the nucleus of an infected cell. Capsid stability is linked to the presence of cofactors, including inositol hexakisphosphate (IP_6_) that bind to pores found in the capsid. Using extensive all-atom molecular dynamics simulations of HIV-1 cores imaged from cryo-electron tomography (cryo-ET) in intact virions, which contain IP_6_ and a ribonucleoprotein complex, we find markedly striated patterns of strain on capsid lattices. The presence of these cofactors also increases rigidity of the capsid. Conformational analysis of capsid (CA) proteins show CA accommodates strain by locally flexing away from structures resolved using x-ray crystallography and cryo-electron microscopy. Then, cryo-ET of HIV-1 cores undergoing endogenous reverse transcription demonstrate that lattice strain increases in the capsid prior to mechanical failure and that the capsid ruptures by crack propagation along regions of high strain. These results uncover HIV-1 capsid properties involved in their critical disassembly process.

**Significance statement:** The mature capsids of HIV-1 are transiently stable complexes that self-assemble around the viral genome during maturation, and uncoat to release preintegration complexes that archive a double-stranded DNA copy of the virus in the host cell genome. However, a detailed view of how HIV cores rupture remains lacking. Here, we elucidate the physical properties involved in capsid rupture using a combination of large-scale all-atom molecular dynamics simulations and cryo-electron tomography. We find that intrinsic strain on the capsid forms highly correlated patterns along the capsid surface, along which cracks propagate. Capsid rigidity also increases with high strain. Our findings provide fundamental insight into viral capsid uncoating.

## Introduction

The mature capsids of HIV-1 are large fullerene-like protein complexes that are comprised of more than a thousand copies of the capsid protein (CA) (1). During replication, viral particles that bud from host cells are initially immature and composed primarily of a spherical array of Gag proteins that link essential viral proteins and enzymes into a linear polypeptide chain (2). As the virus matures extracellularly, proteolytic cleavage of Gag releases the capsid protein, which self-assembles in a cone-shaped geometry and packages two copies of the retroviral genome and associated enzymes into the capsid. Mature capsids are deposited during the fusion of HIV particles with the plasma membrane, in which the lipid envelope and embedded proteins are lost, leaving behind the core of the HIV-1 virus i.e., the capsid and contents, in the cytoplasm of cells.

Capsids play essential roles during replication, by transporting viral genetic material deep into the host cell (3). Pores in the capsid can bind or import small molecules, including inositol phosphates (IP_6_) and nucleotides (dNTPs) (4, 5), and the binding of IP_6_ increases the stable lifetimes of the capsid and promotes the assembly of CA into fullerene structures (6, 7). Cryo-electron tomography (cryo-ET) and other techniques have recently demonstrated that viral cores are imported with apparently intact capsids across the nuclear pore of infected cells (8–11). Reverse transcription processes inside the capsid can rupture the core as seen in both atomic-force microscopy and cryo-ET experiments (12, 13) owing to an increased internal pressure on the capsid during the conversion of RNA to DNA. Uncoating of the capsid is a critical replication event, releasing enzymes and nucleic acids that integrate a copy of the virus in the host genome. Yet, little is known about the physical properties underlying capsid rupture.

To investigate the structural and mechanical properties of HIV-1 capsids that lead to rupture and disassembly, we have performed large-scale all-atom molecular dynamics (AAMD) simulations of HIV-1 capsids, containing native cofactors including IP_6_, and a ribonucleoprotein complex (RNP). Analysis of the strain induced on the capsid, reveals spatially correlated patterns, indicating that the capsid collectively cracks open along regions of high strain rather than slowly disassembles. Local fluctuations in the capsid volume decrease concomitantly with increased strain, consistent with a mechanical rigidification of the capsid in response to IP_6_ and the RNP. Calculated free energy landscapes also reveal shifts in the conformational ensembles of CA in response to increased strain. Cryo-ET imaging of *in vitro* reconstituted HIV-1 cores incubated with nucleotides then add further insight into the temporal sequence of events during disassembly to demonstrate that strain is maximal prior to the formation of cracks in the capsid. These results show that capsids are intrinsically strained and help to characterize the molecular processes by which viral cores rupture.

## Results

Our AAMD simulations of HIV-1 cores contained a total simulation size ranging from 44 to 76 million atoms. In prior AA MD simulations (14, 15), a model for the empty capsid shell, enclosing only water with neither nucleic acid contents nor the binding of inositol phosphates (i.e., IP_6_) was constructed from low-resolution cryo-ET. In contrast, we derived six atomic models from fullerene lattice maps of the complete capsid, with the positions and orientations of lattice components determined by an iterative alignment based on tomograms of intact HIV-1 virions (Fig. 1A) (16, 17). IP_6_ polyanions were added to each CA hexamer or pentamer pore at the binding site, a location 2.7 Å above the R18 ring (5, 7). In the absence of atomic resolution into the structure of the ribonucleoprotein complex (RNP), we constructed a model of two copies of the 9-kilobase pair RNA genome in complex with nucleocapsid proteins consistent with experimental secondary structure constraints from selective 2’-hydroxyl acylation analyzed by primer extension (SHAPE) analysis (18) to mimic the HIV-1 RNP. Core particles were simulated at successively increasing levels of detail ranging from capsid shells containing only water to more realistic capsids containing the RNP model and IP_6_ molecules (Fig. 1 B to E) (see Materials and Methods for a complete description). The Cα positions of the atomic model for the capsids containing the RNP and IP_6_ overlapped well with the 6.8 Å cryo–ET density of the CA hexamer and 8.8 Å cryo-ET density of the CA pentamer positioned in the lattice map (Fig. S1). In aggregate, the AA MD simulations totaled 1.6 μs across the HIV-1 core particles (table S1).

**Figure 1.**
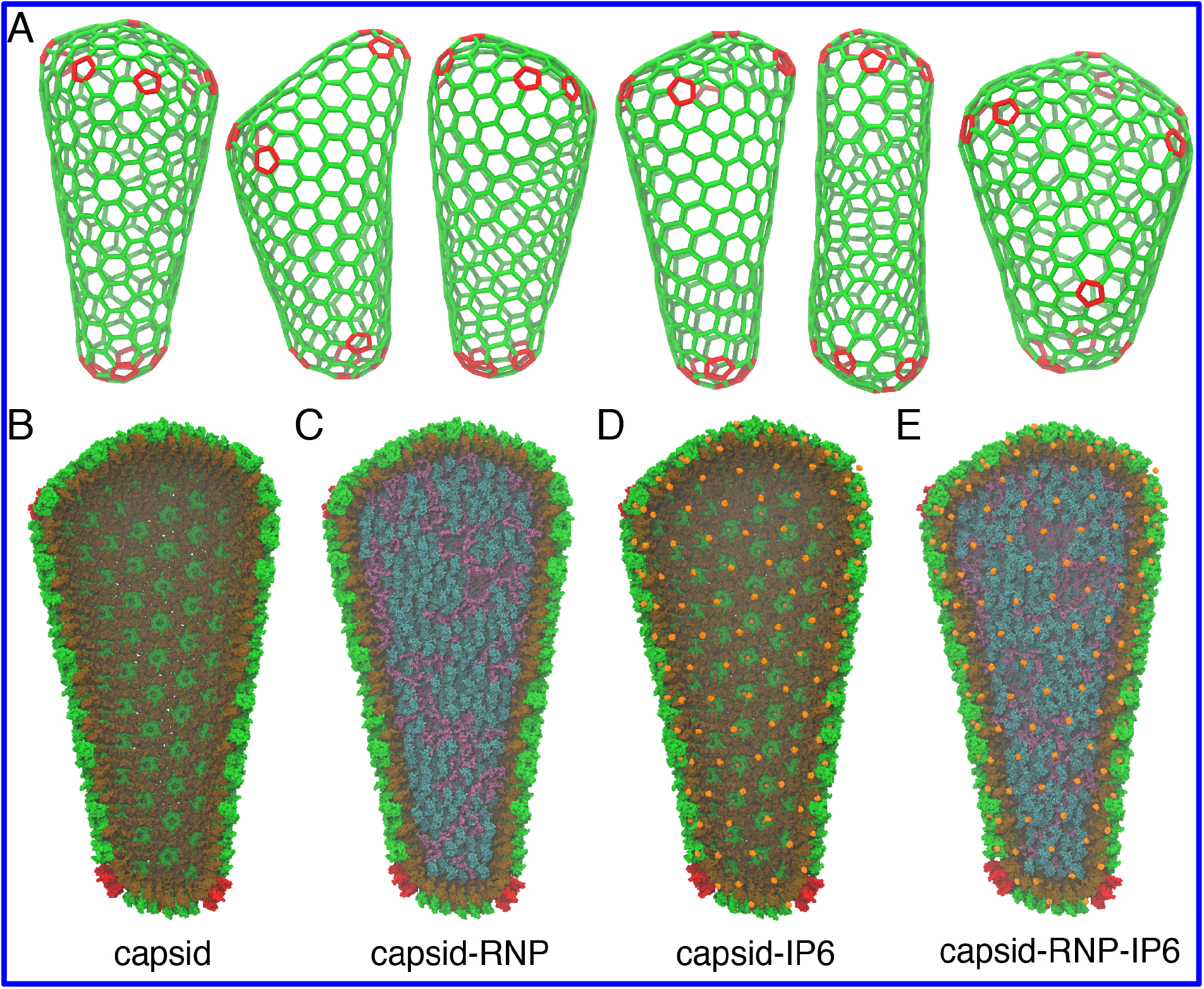
Mature HIV-1 capsids are pleomorphic. **(A)** Fullerene geometries for the HIV-1 capsid were derived from cryo-ET images of intact virions (16). Atomic models for the capsid contain either **(B)** liquid water in the capsid interior, **(C)** a ribonucleoprotein (RNP) complex model, **(D)** inositol hexakisphosphate (IP_6_) molecules bound to the capsid pores, or **(E)** both the RNP and IP_6_. The CA amino terminal domain (NTD) and carboxyl terminal domain (CTD) are colored in green and brown, whereas genomic RNA, nucleocapsid proteins, and IP_6_ molecules are in purple, teal, and orange, respectively. Pentamer defects are colored in red.

### Capsid Strain and Rigidity

During the AAMD simulations, the viral capsids remained intact. Empty capsids in bulk solvent without IP_6_ did not dissociate nor reassemble into more stable helical or spherical arrangements within the timescales simulated. Local deformations in materials and proteins have elucidated the mechanical properties of materials under stress and conformational changes resulting from protein–ligand interactions (19). To quantify these features, we computed per-particle strain tensors (***ε***), for the center-of-masses of five amino acid residue segments in the CA domain. The volumetric strain 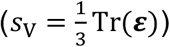 measures the propensity for a particular region of the capsid to either swell or shrink (Fig. 2A). Empty cores containing liquid water had relatively little strain (CA; ⟨|*s*_V_|⟩ = 2.4 × 10^−2^), which was distributed randomly across the capsid. Strain increased In the presence of the RNP complex (CA-RNP; ⟨|*s*_V_|⟩ = 3.3 × 10^−2^). IP_6_ binding induced even larger effects that were also more spatially correlated than the capsid containing just the RNP (CA-IP_6_; ⟨|*s*_V_|⟩ = 3.6 × 10^−2^). Cores containing an RNP complex and IP_6_ molecules were the most strained with patterns that formed unexpected striations (Fig. 2A) along the capsid surface (CA-RNP-IP_6_; ⟨|*s*_V_|⟩ = 5.1 × 10^−2^). Different capsid structures showed a similar trend of increased strain in the presence of IP_6_ and RNP, but with slightly altered strain patterns that are attributable to differences between core morphologies (Fig. S2).

**Figure 2.**
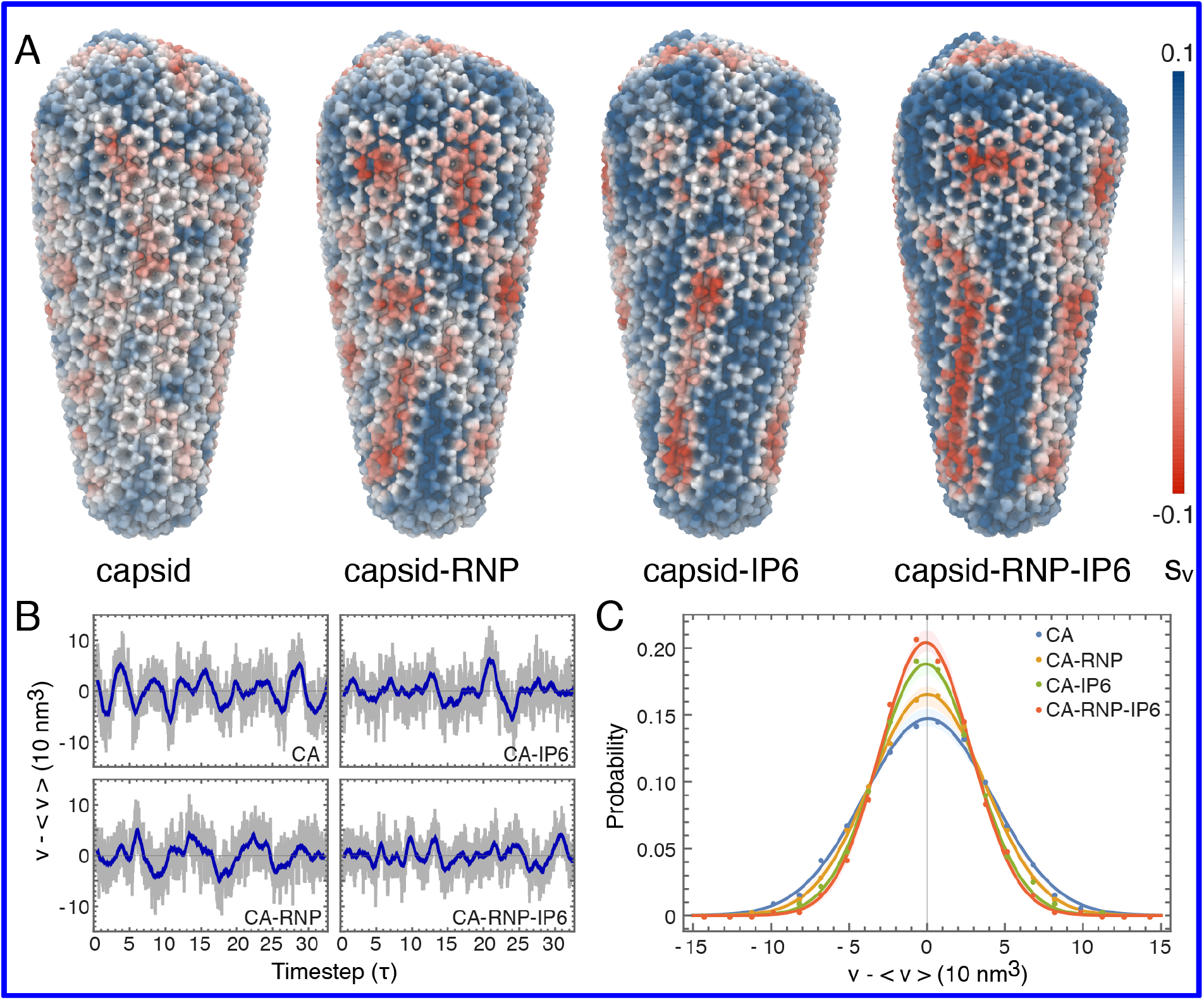
Strain and fluctuation analysis of the HIV-1 capsid. **(A)** The first invariant of the strain tensor (*s*_V_) or volumetric strain was computed for each capsid core complex. Red and blue colors correspond to compressive and expansive strain, respectively. **(B)** Fluctuations in the HIV-1 core volumes, measured as deviations from the average capsid volume (⟨*V*⟩ = 1.45 × 10^5^ nm^3^). The subsampled timestep, τ, is 4 ns. Gray colors denote the instantaneous volume, whereas the blue line denotes a moving average within a one-timestep window. **(C)** Probability distributions for the volume fluctuation amplitudes in each capsid. Closed circles correspond to measured data, while the solid line indicates a Gaussian fit. The shaded regions denote the error in the distribution determined by block averaging.

To assess whether capsid properties change in response to IP_6_ or RNP, we monitored the internal volumes of each core during the simulations. Although each pleomorphic capsid differed in size (Fig. 1 A), average core volumes ranged from 1.15– 1.53 × 10^5^ nm^3^, with fluctuations of several tens of cubic nanometers about the equilibrium. Mean-free volume fluctuations for the largest core are shown in Fig. 2B. Fluctuation amplitudes decreased markedly in the presence of either IP_6_ or the RNP, consistent with higher core rigidity (Fig. 2C, CA-IP_6_ vs. CA-RNP) (std. dev.: σ_CA_= 41.1 nm^3^; σ_CA-RNP_= 36.3 nm^3^; σ_CA-IP6_= 31.2 nm^3^; σ_CA-RNP-IP6_= 29.2 nm^3^). Decreases in the volume fluctuation amplitudes were consistent across cores of different morphologies (Fig. S3). Fourier analysis showed a shift in the peak-to-peak frequencies of the dominant mode towards lower frequencies with the RNP, indicating that the capsids fluctuate more slowly (CA: *ν* = 57.8 MHz; CA-RNP: *ν* = 39.4 MHz) (Fig. S4). As the core rigidifies further upon IP_6_ binding, the dominant low frequency mode broadens, and the fluctuations are distributed to higher frequency modes, consistent with a stiffer capsid. Negatively charged IP_6_ molecules bind tightly to an arginine ring in pores distributed throughout the capsid (4, 5, 7). RNP interactions with CA, on the other hand, could occur through interactions of positively charged residues on the flexible CTD tail of CA with the RNP (7). The cofactor interactions at the pore and CTD tail change the conformational flexibility of CA resulting in an increase in capsid rigidity that introduces strain on the underlying lattice.

### Conformational Analysis of CA Proteins

To examine the conformations CA proteins adopt in the actual capsid, we computed a 3D free energy landscape (Fig. 3 A,B) from the spatial distribution occupied by the non-hydrogen atoms of CA across the pleomorphic lattice for the capsids containing IP_6_ and RNP. At high contours (Δ*G* = 0.5 kcal/mol), density for the protein backbone of CA is clearly visible, whereas at low contours (Δ*G* = 4.4 kcal/mol), variability in the ensemble of CA structures undergoing dynamical motion results in larger volumes (Fig. 3B). Less structured regions of the protein, including the CTD tail, β hairpin, and CypA binding loop, had higher variability, and were more mobile. The hinge connecting the CA NTD and CTD was more ordered than the other unstructured regions, possibly owing to NTD– NTD and CTD–CTD contacts in the adjacent CA domains of the capsid. The conformations of individual CA monomers in pentamers and hexamers resolved by cryo-ET (16) show small differences in the relative orientations of the NTD and CTD (Fig 3C). Relative free energy differences between the conformations were quite small (Δ*G* < 0.2 kcal/mol), indicating that CA can switch between the two states in the capsid under thermal motion.

**Figure 3.**
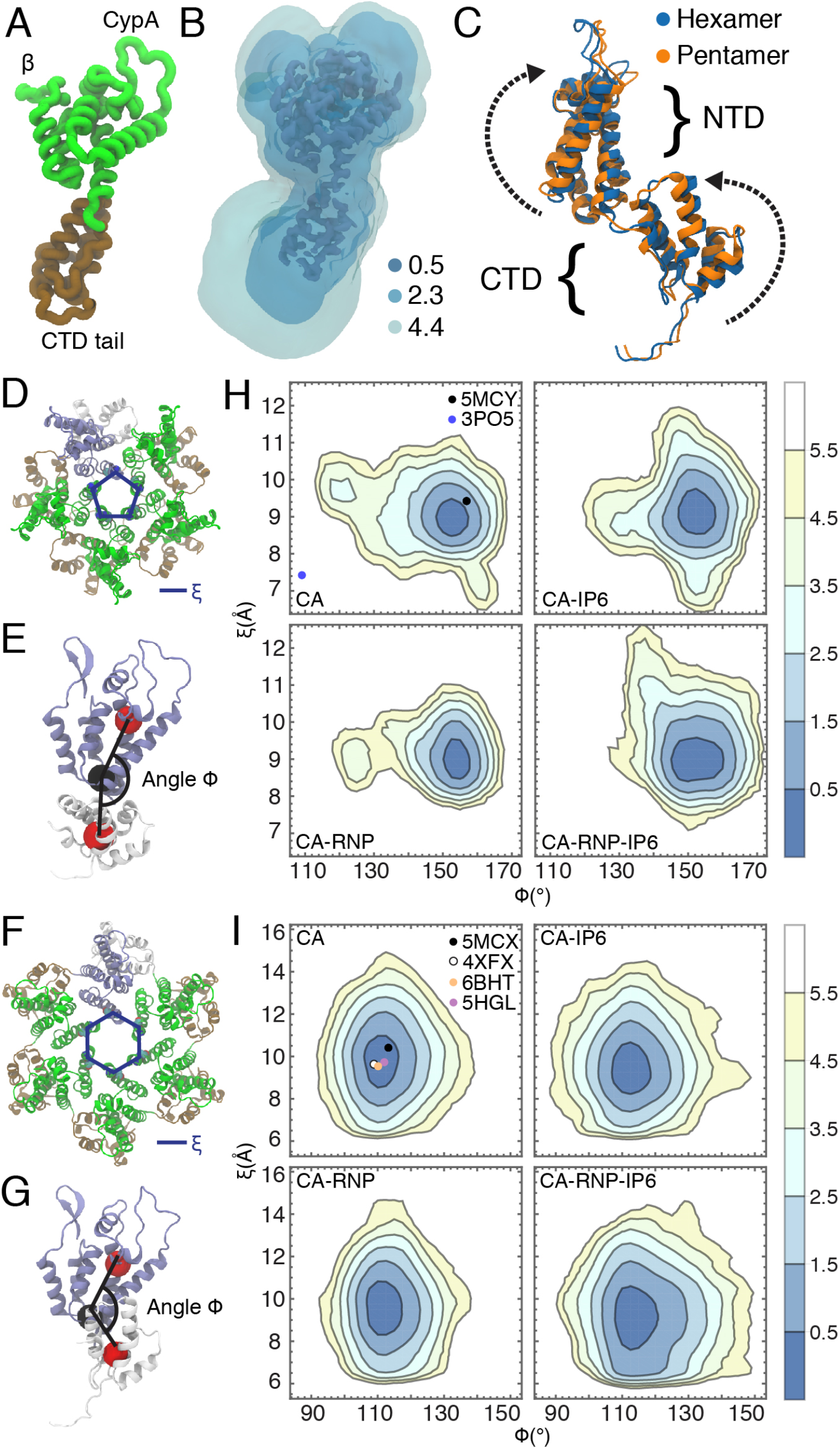
Conformational analysis of CA domain proteins. **(A)** A monomer of the CA protein. **(B)** The 3D potential of mean force (PMF) for CA conformations contoured at 0.5, 2.3 and 4.4 kcal/mol with respect to the Cartesian coordinates of the protein heavy atoms. **(C)** Differences in CA monomer conformations between the cryo-EM structure of the hexamer (PDB ID: 5MCX) and cryo-EM structure of the pentamer (PDB ID: 5MCY) consist of rotations in the NTD and CTD. The parameter, ξ, is used to describe the pore size and is defined as the center-of-mass distance between residues N21 and A22. The angle Φ is used to describe the relative orientation of a NTD and adjacent CTD and is defined as the angle between three center-of-masses across the NTD and CTD (red and black spheres). ξ and Φ are shown for the CA hexamer **(D, E**) and pentamer **(F**,**G). (H)** (ξ, Φ) distributions for the hexamers in each capsid. **(I)** (ξ, Φ) distributions for the pentamers in each capsid. Contour lines correspond to increments of *f* = − log *p*, where *p* is the probability. Closed circles correspond to (ξ, Φ) values from experimental structures in the Protein Data Bank.

Interfacial contacts between the NTD of one CA domain and the CTD of the adjacent CA domain differed appreciably in the pentamer and hexamer (Fig. 3D to I). We employed an angle parameter, *ϕ*, defined as the angle between amino acid backbone center-of-masses in the NTD helices and CTD helices, using the base of the NTD helices as a pivot, to assess the relative orientation of the NTD and adjacent CTD across hexamers and pentamers in the capsid during MD (Fig. 3E, G). The angle, *ϕ*, was larger in the cryo-ET structure (16) of the pentamer (*ϕ* = 157°) than in the hexamer (*ϕ* = 113°) reflecting a greater curvature of pentamers over hexamers. Pore sizes were also monitored using a parameter, *ξ*, defined as the distance between the center-of-masses of N21 and A22 on adjacent CA monomers (Fig. 3D, F). Pentamers in flexible, empty capsids had high curvatures and small pore sizes (*ξ*_CA_, *ϕ*_CA_) = (7– 11 Å, 115– 167°), although a subpopulation (*f* = 2.5– 3.5) shifted to a lower curvature closer to that of the x-ray crystal structure of a flat pentamer (20) with engineered disulfide crosslinks between N21 and A22 (PDB ID: 3P05) (*ξ, ϕ*) = (7.5 Å, 107°) (Fig. 3H). IP_6_ binding led to a conformational shift in pentamer distributions towards larger pore sizes (*ξ*_RNP–IP6_ = 7– 12.5 Å), and capsids with both IP_6_ binding and the RNP increased pentamer curvature (*ϕ*_RNP_ = 119 − 168°, *ϕ*_IP6_ = 121 − 170°, *ϕ*_RNP–IP6_ = 128 − 171°). CA hexamers occupied wider pores, but flatter curvatures (lower *ϕ*), (*ξ*_CA_, *ϕ*_CA_) = (6.5– 15.6 Å, 92– 136°) compared to pentamers. IP_6_ binding and the RNP did not significantly affect pore size in hexamers, although there was a greater variation in hexamer curvature (*ϕ*_IP6_ = 90 − 147°, *ϕ*_RNP–IP6_ = 91 − 150°), owing to the strain induced on the overall capsid lattice (Fig 3I). X-ray crystal and cryo-EM structures agree with the minima of both curvature and pore size distributions, and resided within the lowest two contours of the (*ξ, ϕ*) distributions (*f* < 1.5), but CA conformations deviate away from these minima under increasing lattice strain.

The NTD-CTD interface is the binding site for FG motifs required for nuclear import of the capsid (21) and small-molecule inhibitors including PF74 (22) and Lenacapavir (23) that disrupt HIV function. Our simulations indicate these binding pockets are modulated by the presence of IP_6_ and the RNP. CA hexamers are less perturbed than pentamers, but exhibited conformational shifts towards increased *ϕ* angles and larger binding pockets. Pentamers on the other hand, existed in two dynamic subpopulations – one of which was consistent with x-ray structure of a flat pentamer, and another consistent with a curved pentamer found in intact virions. The presence of strain (e.g., by IP_6_ and RNP) induced local changes in the CA lattice that favored highly curved pentamers, and a more open binding site at the NTD-CTD interface.

### Mechanical Rupture of the HIV-1 Capsid

Coarse-grained (CG) simulations in which the internal pressure on the capsid was varied were performed to examine how CA-CA interaction strengths altered capsid disassembly behavior (Fig. 4). To uniformly increase the pressure inside the capsid, particles with excluded volume interactions were added in the capsid interior, until the point of rupture. Below a critical degree of CA-CA interaction (*k* = 1.20 kcal/mol), the capsid disassembles even without any internal pressure (*P* = 0), whereas above this interaction strength the capsid remains stable, indicating that there is a minimum lattice energy required to stabilize the fullerene geometry of the CA lattice. The pressure required for rupture increased with increasing CA-CA interaction strength. At low CA-CA interaction strengths (*k* < 1.30 kcal/mol), small separations in the lattice lead to the gradual dissociation of CA and disassembly of the capsid along the separations. In the CG simulations, pentamers dissociated more quickly from the lattice, owing to the fewer attractive contacts the pentamers had in the lattice compared to hexamers. At higher CA-CA interaction strengths (*k* ≥ 1.30 kcal/mol), rupture of the capsid was more collective, with larger cracks that formed across the capsid surface. These CG simulations highlight that the intermediates formed during rupture of the capsids are sensitive to the strength or rigidity of CA-CA interactions in the capsid.

**Figure 4.**
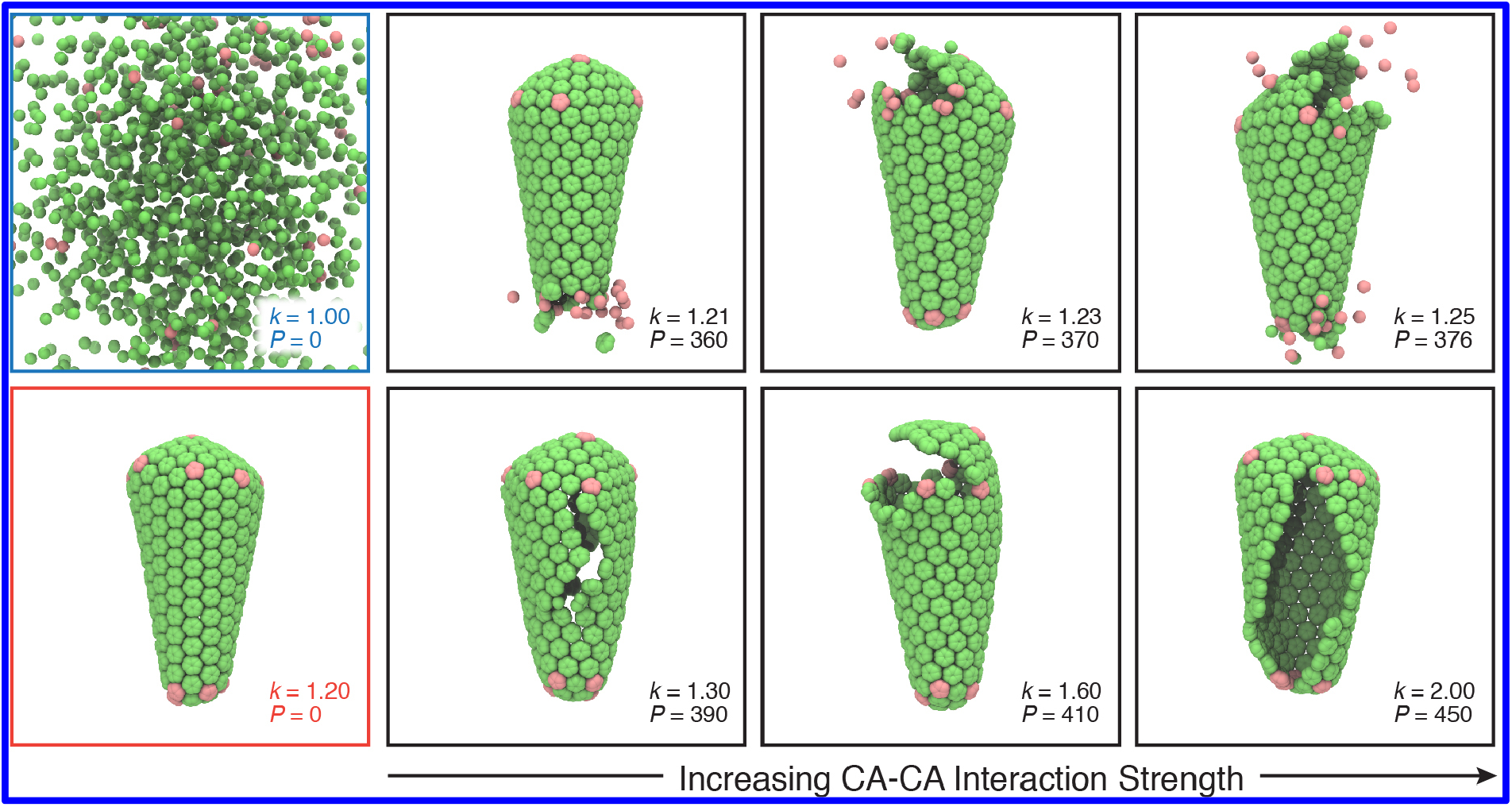
Rupture of the capsid in CG simulations. A series of snapshots show the mechanical failure of the capsid as internal pressure is increased until the point of rupture, taken from the trajectories at time *τ* = 10^6^ CG MD timesteps. The strength of CA-CA interactions (*k*) is given in units of kcal/mol, whereas the pressure (*P*) is given in units of spherical particles that are added to the capsid interior. At *k* = 1.00 kcal/mol; *P* = 0, the capsid slowly disassembles (blue panel). At *k* = 1.20; *P* = 0, the fullerene lattice remains stable (red panel). With low CA-CA interaction strengths (*k* = 1.21, 1.23, 1.25 kcal/mol), small lattice separations lead to a gradual disassembly of the capsid. With high CA-CA interaction strengths (*k* = 1.30, 1.60, 2.00 kcal/mol), rupture is characterized by the formation of larger cracks in the capsid.

### Core Rupture During Endogenous Reverse Transcription

Cryo-ET imaging and lattice mapping were used to probe how core structure changed during reverse transcription (13). In brief, HIV-1 cores were released from purified virions by permeabilizing the viral membranes with the pore-forming melittin peptide and then stabilized by the addition of IP_6_ at native cellular concentrations. Reverse transcription was initiated by adding dNTPs at concentrations found in CD4^+^ T cell cytoplasms (24). After four hours of incubation at 37 °C, a total of 14 cores were found in the tomograms and mapped. Capsid structures showed a range of intermediates from intact cones to partially cracked capsids to nearly completely disassembled (Fig. 5A, images 1-14). We examined lattice separations in each core using a local order parameter, *χ*, which measures the near-neighbor contacts of a particle. In our AAMD simulations, lattice separations correlated with the strain, as expansive strain caused slight, but measurable deviations in lattice separation (*χ* < 1.0), whereas compressive strain induced more closely packed lattices (*χ* > 1.0) (Fig. 5B). No neighboring particles are present at low values of *χ*, which provided a quantitative metric for the degree of cracking across the core (*χ* < 0.55). Interestingly, tubular “pill” shaped capsids (Fig. 5A, 1) were nearly perfect fullerenes with no cracks present, whereas other more conical capsids had cracks that were not visually obvious prior to computing *χ* in initial inspections (Fig. 5C). The various defects found in the experimental capsid structures included small ruptures along the narrow and broad end of the fullerene cone as well as large cracks that formed along the length of the capsid (Fig. S5). Intermediate values for *χ* correspond to small deviations and separations that reflect the expansive strain on the capsid (0.75 < *χ* < 0.9). Analysis of *χ* at these intermediate values revealed a bimodal distribution (Fig 5 D). We observed largely intact capsids with minimal strain (Fig 5A 1,2) and high strain (Fig. 5A 3,4). Small ruptures or cracks in the lattice decrease the strain (Fig 5A 5,6,7,8), which then increase during further stages of disassembly (Fig 5A 9, 10, 11). We interpret these results to indicate that mechanical failure of the capsid and loss of capsid integrity lowers the strain, which then increases as the pre-integration complex exits the core, and that reverse-transcribing capsids are strained prior to and during rupture of the capsid.

**Figure 5.**
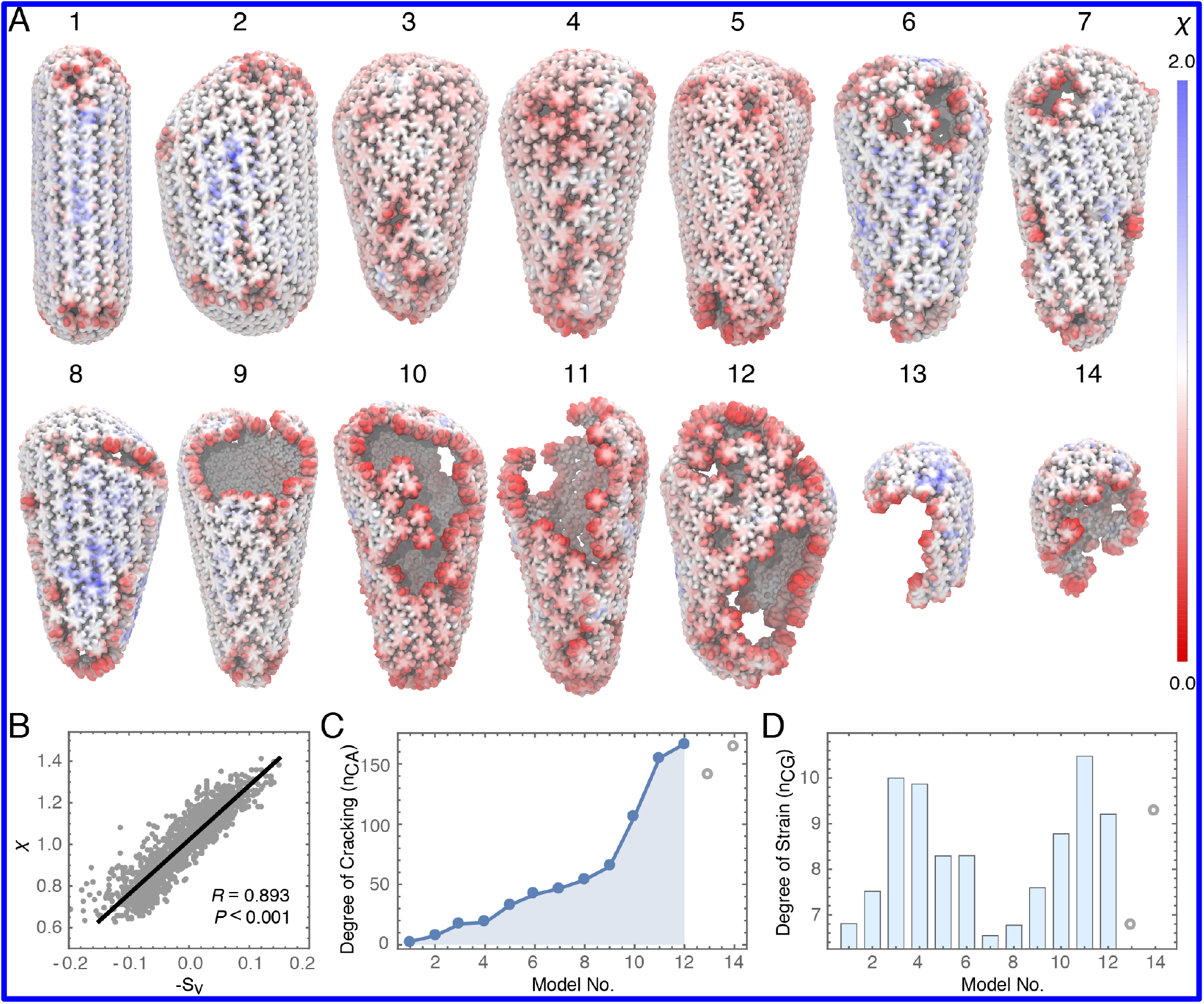
Cryo-ET of HIV-1 cores during rupture. **(A)** Cores were imaged by cryo-ET and lattice mapping during endogenous reverse transcription. Each structure (images 1-14) is colored by a local order parameter (*χ*) that quantifies the degree of separation in the lattice. An ideally spaced particle in the lattice has a separation of *χ* = 1.0. Red colors correspond to particles further apart on average from near-neighbors, whereas blue colors correspond to particles closer to near-neighbors. **(B)** Correlation between the strain (*s*_V_) and lattice separation (*χ*) measured in the MD simulations. **(C)** The degree of cracking in each capsid model measured by the number of CA monomers containing a particle with *χ* < 0.55. **(D)** The degree of strain in each capsid model measured by the number of particles (10^3^) with 0.75 < *χ* < 0.9. Capsids 1–14 in panel A were ordered from no fractures to completely ruptured. Outliers in which the capsid was mostly disassembled are shown in open circles.

## Discussion

We simulated entire HIV-1 core particles derived from cryo-ET at an atomic level of detail and computed the intrinsic strain on the capsid induced by IP_6_ and RNP. Unexpected correlated strain patterns formed on the capsid surface, and analysis of core volume fluctuations showed that the capsid mechanically rigidifies in response to the presence of these co-factors. Our conformational analysis of CA proteins in the capsid also indicates that pentamers and hexamers have a surprising degree of dynamic flexibility that changes under strain. The CA pentamer adopts two distinct states, one corresponding to a state with flat curvature and another corresponding to a curved state with a shift towards more highly curved conformations under increased strain. CG MD of the capsid under pressure demonstrates that rupture is initiated by small separations in the CA lattice that lead to disassembly of the capsid at regions of high strain, and as the CA-CA interaction strength is increased in the model the rupturing becomes more collective with large cracks forming in the capsid. Cryo-ET of core rupture during endogenous reverse transcription reveals that both small separations and large cracks in the capsid are present, and show that lattice strain is locally maximal prior to mechanical failure. The capsids rupture in a fashion consistent with the propagation of cracks along highly strained regions, as observed in the simulations.

In biochemical assays (13), overstabilization at high IP_6_ concentrations results in cores that do not produce reverse transcription products, and the binding of small-molecule inhibitors such as Lenacapavir (23, 25) can weaken capsid integrity and accelerate fracturing of the capsid (13), suggesting HIV-1 cores have physical properties that can be altered to disrupt retroviral life cycle processes. During trafficking from the cytoplasm to the nucleus of infected cells, viral capsids are exposed to interactions with a variety of host proteins. Cyclophilin A binds to and enhance stability of the capsid in a concentration dependent fashion (26) in the cytoplasm. Nucleoporins bind to and import viral cores across the nuclear pore of host cells (27, 28). The cleavage and polyadenylation specific factor 6 (CPSF6) interacts with capsid complexes (22) and is involved in nuclear import and trafficking to integration sites (29, 30). Could these interactions with host proteins along with increasing internal pressure from reverse transcription weaken capsid integrity and rupture the capsid? Additional investigations should therefore further probe the molecular mechanisms involved in viral core interactions within the host cell.

## Materials and Methods

### All-atoms models of the capsid core

Initial atomic models for the CA hexamer and pentamer were constructed from the cryo-ET structure of the CA hexamer (PDB ID: 5MCX) and CA pentamer (PDB ID: 5MCY) derived from intact virus particles. Amino acid side chain conformations were modeled based on the x-ray crystal structure of the CA hexamer (PDB ID: 4XFX) and disulfide-crosslinked CA pentamer (PDB ID: 3P05). Missing protein backbones residues were built using MODELLER (31), and missing side chains were built using SCWRL4 (32). C-terminal domain (CTD) tails for CA were transplanted on to the model using the NMR structure for the tail (residues: 220–231) (PDB ID: 2KOD) onto the model. Atomic models for the CA hexamer and pentamer were then coarse-grained (CG) at a resolution of one amino acid residue per CG particle. Each CG hexamer and pentamer subunit was constrained as a rigid body, and positioned at the Cartesian coordinates and Euler angles that maximized overlap between the atomic model and the cryo-ET lattice map derived from intact virions (16).

CG models of the capsids were briefly relaxed in a 20 ps Langevin dynamics run under the canonical (constant NVT) ensemble with the large-scale atomic/molecular massively parallel simulator (LAMMPS) (33). To maintain overall protein shape, each CG particle included excluded volume interactions using a soft cosine potential *U*_Vol_ = *A* [1 + cos (*π r*/*r*_*c*_)], when *r* < *r*_*c*_. The cutoff (*r*_*c*_) is the onset of excluded volume repulsion between CG particles, and was set to the separation distances found in the CTD interfaces if the distances were less than 10 Å, or otherwise 10 Å. The parameter *A* was taken to be *A* = 100 kcal mol^−1^. The mass of the protein was evenly distributed among the CG beads. CTD contacts were maintained using an elastic network model (ENM) that connected the two groups of CG particles in neighboring CTD dimers (residues: 178–194) in a cutoff distance, *r*_cut_, with harmonic bonds; the potential energy of each bond is: *U*_bond_ = *K*_bond_ (*r* − *r*_0_)^2^, where *r* is the separation distance and *r*_0_ is the equilibrium bond length, set to the distance found in the crystal structure (*r*_cut_ = 10 Å, *K*_bond_ = 0.1 kcal mol^−1^ Å^−2^). Temperature was mainted at 300K with a Langevin thermostat with a damping constant (*t*_damp_ = 5 ps). All-atom (AA) models for each CA hexamer and pentamer were then aligned with each CG capsomere subunit to construct an initial AA model for the capsid. Six complete capsids were constructed corresponding to the fullerene geometries derived from the cryo-ET structures, two of which were selected for additional modeling.

For each of the capsid systems selected, IP_6_ molecules were placed at positions corresponding to the bound x-ray crystal structure, 3.5 Å above an R18 ring that lines the pore of the CA hexamer and pentamer. Approximately 200 IP_6_ molecules were used for each system, corresponding to a single IP_6_ molecule for each capsomere subunit in the fullerene structure. A model of the ribonucleoprotein core was generated using a simplified CG model of RNA. The full length HIV-1 genomic RNA sequence was used (18), and CG particles were added to the capsid interior at a resolution of four beads per nucleotide base pair. The CG particles were labeled corresponding to the atom types in each base pair (adenine: C3’ C8 N6 C2; cytosine: C3’ C6 O2 N4; guanosine: C3’ C8 O6 N2; uracil: C3’ C6 O2 O4) to generate the viral RNA. Bonded interactions were added using harmonic restraints between the neighboring nucleotides to (*K*_bond_ = 1.0 kcal mol^−1^ Å^−2^) and set to the average equilibrium distances between CG particles found between the represented atom types in the x-ray crystallographic structure of a model RNA template (PDB ID: 4GXY) to maintain geometric shape. Secondary structure constraints and base pairing interactions were implemented using harmonic restraints between the nearest-neighbor non-bonded CG particles between base pairs on the basis of high-throughput selective 2-hydroxyl acylation analyzed by primer extension (SHAPE) reactivity data (18) (*K*_SS_ = 0.1 kcal mol^−1^ Å^−2^). Two copies of the 9 kilobase pair genome were modeled and positioned in the interior of the capsid. CG particles for the nucleocapsid proteins (NC) were mixed with the RNA at random positions in the capsid interior, until there was a 1:1 correspondence between the number of CA and NC domains for the RNP. The CG RNP model was energy minimized and relaxed under a brief 100 ps Langevin dynamics run. All-atom models for the RNA and nucleocapsid proteins (PDB ID: 1A1T) were then fit to the CG RNP, and subsequently energy minimized and equilibrated as described below.

### All-atom MD simulations of viral capsid cores

Solvated capsids were of sizes ranging from 44–76 million atoms including water molecules and ions. Na^+^ and Cl^-^ ions were added to the bulk solution until the salt concentration was 150 mM NaCl to produce an electrostatically neutral system. Periodic boundary conditions were imposed on an orthorhombic unit cell ranging from 55.2 nm × 58.9 nm × 128.5 nm – 72.8 nm × 76.3 nm × 141.9 nm, and contained a solvent buffer of 10 nm in the (x, y, z) dimensions away from nonsolvent atoms. The AA potential energy function CHARMM36m (34, 35) for proteins and the TIP3P (36) potential energy function for water were used. The AA systems were energy minimized and equilibrated under constant pressure and temperature (NPT) conditions. Simulations in the constant NPT ensemble were performed using a Langevin thermostat at 310 K and a Nosé-Hoover Langevin barostat at 1 atm. Bond-lengths for hydrogen atoms were constrained using the SHAKE algorithm (37). An r-RESPA integrator was used with a timestep of 2 fs; long-range electrostatics were computed every 4 fs (38). Long-range electrostatics were calculated using the particle mesh Ewald algorithm, and short-ranged, nonbonded interactions were truncated at 12 Å (39). IP_6_ molecules were parameterized using the CHARMM General Force Field (CGenFF) (40). All simulations used the AAMD simulation package NAMD 2.14 (41). Production-level runs were performed on six capsid containing liquid water and two capsids containing IP_6_, the RNP core, or both IP_6_ and RNP. For all subsequent analysis AAMD trajectories were sampled at 0.04-ns intervals.

### Strain calculations

The center-of-masses for every five amino acid residues in the CA domains were used to subsample the capsid structure. For each center-of-mass, a deformation tensor that describes the local deformation of a point particle in a system relative to it’s neighboring particles was calculated: 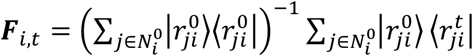, where 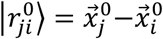 is the difference between the Cartesian coordinates of neighboring particles *j* and *i* with respect to the reference configuration, similarly 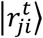 is the difference in coordinates at time *t*, and 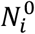 is the set of particles in the local neighborhood of *i* in the reference configuration, within a cutoff distance of 10 nm. Note that the deformation tensor, ***F***_*i,t*_, can be determined by minimizing 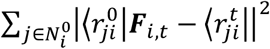.The reference configuration was the fitted atomic model of the fullerene capsid after CG relaxation. The per-particle Green-Lagrange strain tensor is then, 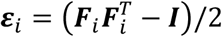, with volumetric and shear (von Mises) strain invariants given by: *s*_V_ = Tr(***ε***)/3 and *s*_s_ = [Tr(***ε***^2^) − Tr(***ε***)^2^/3]^1/2^ respectively. Strain tensors were computed for each frame of the trajectories, until the invariants converged to an equilibrium value (e.g., 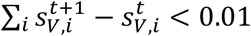) at frame *t* ∼ 32 ns. Strain patterns persisted for the remainder of the simulation time (see also Table S1). Average strain magnitudes, ⟨|*s*_V_|⟩, were measured using the strain corresponding to residue 110.

### Core volume analysis

Core volumes of the capsid were computed using a Monte-Carlo sampling strategy. Cartesian coordinates for the Cα atoms of the CA domains were monitored and used to subsample the capsid structure. For each iteration, a point within the periodic simulation cell were randomly generated 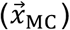, and assigned to the capsid interior, if 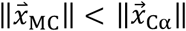, where 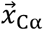 is the position of the Cα atom that maximizes the projection of the point onto the capsid structure 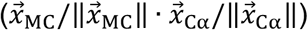. 10^9^ iterations were performed, and core volumes were calculated from the total fraction of points inside the capsid, accumulated for each 0.04 ns interval frame. Fluctuations from the mean volume (*V* − ⟨*V*⟩) were measured in each capsid trajectory. Normalized fluctuation probabilities were obtained by binning the amplitudes with a 15 nm step-size, and subsequently fit to Gaussian distributions. Statistical uncertainty was assessed by block averaging. For each distribution, the time series was divided into three blocks. The standard deviation in fluctuation amplitudes for the three blocks is reported. Using 3-5 blocks, gave qualitatively similar results. Fourier analysis was performed by computing the discrete Fourier transform on the mean-free volume fluctuations (42).

### CA Conformational distributions

AAMD trajectories from production-level simulations of CA systems were subsampled at 0.04-ns intervals and aggregated. CA monomers in the capsid were aligned by minimizing the RMSD of the protein backbone atoms with respect to the crystallographic structure (PDB ID: 4XFX). Cartesian coordinates for the nonhydrogen atoms of CA were recorded, and the density of atomic positions 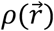 was calculated, using a hard-sphere van der Waals approximation on a discretized grid with a spacing of 0.5 Å × 0.5 Å × 0.5 Å as in (43). The 3D density map for the CA monomer was subsequently contoured at *G* = 0.5, 2.3, and 4.4 kcal/mol, where *G* = −*k*_B_ *T* log *ρ*. The parameter, *ξ*, was used to specify the distance between the center-of-masses of the amino acid backbones of residue 21 and residue 22 on the pore helices (H1) of adjacent CA domains. The angle, *ϕ* is defined as the angle between three center-of-masses in the amino acid backbone, two of which were positioned on the CA NTD (residues 50–52, 78–80, and 128–130; residues 63–65, 144–146, and 58–60) and the third positioned at the adjacent CA CTD (residues 188–192, 166–170, 196–200). (*ξ, ϕ*) values were recorded for each 0.04-ns frame and aggregated across each of the simulated capsid systems (CA, CA-RNP, CA-IP_6_, and CA-RNA-IP_6_). Successive contours for the (*ξ, ϕ*) distributions are given in units of − log *p*, where *p* is the probability.

### CG simulations of capsid rupture

CG models of the capsid were generated from the all-atom models by computing the center-of-mass of each CA monomer at a resolution of 1 CG particle or “bead” per CA monomer. CA-CA interactions were added using a network of Morse bond effective potentials between the 8 nearest-neighbor particles 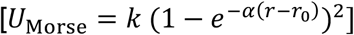, where the equilibrium bond distance, *r*_0_, was set to the distance found between the neighboring center of masses, and the stiffness set to 2 Å^−1^. Excluded volume interactions between CG particles used a soft-cosine potential (*U*_Vol_), with *A* = 100 kcal mol^−1^ and cutoff *r*_*c*_ = 75 Å. All CG MD simulations were performed in constant NVT conditions, with a periodic cell of (2,000 Å × 2,000 Å × 2,000 Å) and a Langevin thermostat at 300 K. CG particles with excluded volume interactions were added every 1,000 CG MD timesteps in a spherical region with radius 150 Å centered at the origin, until the capsid ruptured. All simulations were performed in LAMMPS and proceeded for 10^6^ CG MD timesteps.

### Cryo-ET imaging of fractured cores

Endogenous reverse transcription (ERT) starting with purified HIV-1 virions was performed as described (13). Samples of cores undergoing ERT for 4 h at 37 °C were mixed with equal volumes of 10-nm BSA Gold Tracer (Electron Microscopy Sciences), and 3.5 uL aliquots were applied onto glow-discharged Quantifoil grids (Electron Microscopy Sciences), blotted to near-dryness, and then plunge-frozen into liquid ethane. Cryotomograms were acquired using an FEI Titan Krios electron microscope operating at 300 kV and equipped with either a Falcon III camera or a K3/GIF with a slit width of 20 eV. Tilt series were collected using the data collection software Tomography 4.0 (FEI) with an angular range of −60° to +60°, an angular increment of either 1 or 2°, defocus values of 5 to 10 μm, and a nominal magnification of ×29,000 (Falcon III) or ×33,000 (K3), which correspond to a pixel (px) size of 2.92 (Falcon III) or 2.69 (K3) Å. Tilt series were aligned by using IMOD (44). Weighted back-projection or SIRT was used to reconstruct tomograms in IMOD.

Lattice mapping was performed as described (13) using as a search template a previously-determined cryo-EM map of a HIV-1 CA hexamer (EMD-3465) down sampled to 25 Å. Each capsid surface was modeled and oversampled as a mesh, using surface modeling tools in the Dynamo software package. Between 2000-3000 sub-volumes were exacted from the tomograms, each centered on points on the capsid surface with points distributed as evenly as possible and sampling the entire capsid surface as if it were intact. Multiple iterations of density matching and alignment of the search template with the sub-volumes were performed in Dynamo, which converges to a lattice map containing the positions and orientations of hexamers on the capsid surface. Uncertainty in the lattice map was determined by the cross-correlation between each hexamer and the local density contained in the sub-volumes (Fig. S6). A total of 14 HIV-1 cores were imaged and mapped, and all tomograms containing cores were analyzed.

### Local order parameter (*χ*) analysis

Atomic models for CA hexamers were aligned with the Cartesian coordinates and Euler angles for each subtomogram in the cryo-ET density. For each structure, Cα positions were used to compute a per-particle, order parameter, *χ*, which quantifies the separation in the capsid lattice. *χ* is defined as the number of neighboring particles (within a cutoff distance of 10 nm) of the same chemical identity in a CA monomer (e.g., residue number) *N*_p_, normalized by the average *N*_p_ for a given residue. *χ* values < 1 correspond to lattice separations that were higher than average for the capsid structure, whereas *χ* values > 1 correspond to lattice separations that were lower than average. The capsid with the highest degree of strain (CA-RNP) from the MD simulations was used to measure correlations between *χ* and *s*_V_. The values of *χ* and *s*_V_ were measured for the protein residue with the largest variation in lattice separation (residue 110). The total number of CA monomers that contain a particle with *χ* < 0.55 or *n*_CA_, and the total number of particles with 0.75 < *χ* < 0.9 or *n*_CG_ were measured for each capsid structure.

## Acknowledgements

This research was supported by the National Institute of Allergy and Infectious Diseases (NIAID) of the National Institutes of Health under grant R01-AI154092 (G.A.V.), grant P50-AI150464 for the Center for the Structural Biology of Cellular Host Elements in Egress, Trafficking, and Assembly of HIV (B.K.G.-P., O.P.), and the Medical Research Council as part of UK Research and Innovation (MC_UP_1201/16 to J.A.G.B.). B.K.G.-P. and O.P. were also supported by NIH-NIAID grants R01-AI129678 and R01-AI150479. Computational resources were provided by Frontera at the Texas Advanced Computer Center funded by the National Science Foundation (OAC-1818253). A.Y. gratefully acknowledges postdoctoral fellowship support from NIH-NIAID under grant F32-AI150208. The authors thank Simone Mattei for preparing and sharing cryo-ET data on HIV-1 capsids.

