## Supplemental File for "Strain and rupture of HIV-1 capsids during uncoating"

**This PDF file includes:**

Figures S1 to S6
Table S1

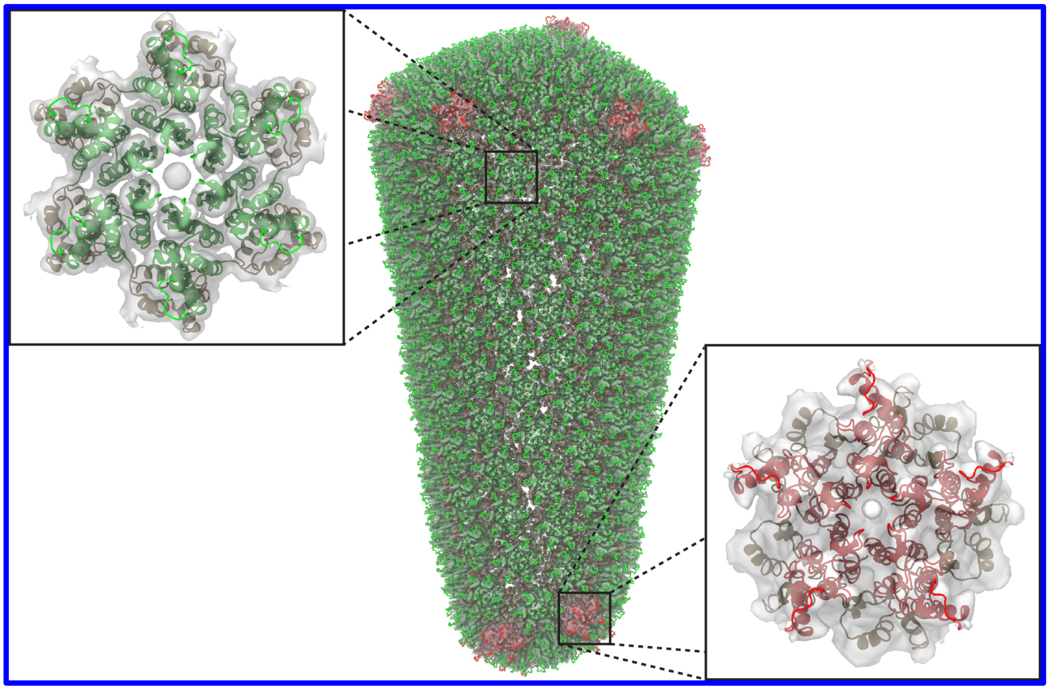

**Figure S1**. **Comparison between the experimental lattice map of the HIV-1 core and the all-atom capsid model with RNP and $IP_6$.** The cryo-ET density for a CA hexamer resolved at 6.8 Å (EMD-3465) and the cryo-ET density of a CA pentamer resolved at 8.8 Å (EMD-3466) were positioned into the lattice map by translating and rotating the two densities to the specified Cartesian coordinates and Euler angles. The densities are overlaid with the Cα positions of the all-atom model for CA-RNP-$IP_6$ (Fig. 1E). (*Inset*, *left*) Expanded view of the all-atom CA hexamer model and EMD-3465. (*Inset*, *right*) Expanded view of the all-atom CA pentamer model and EMD-3466.

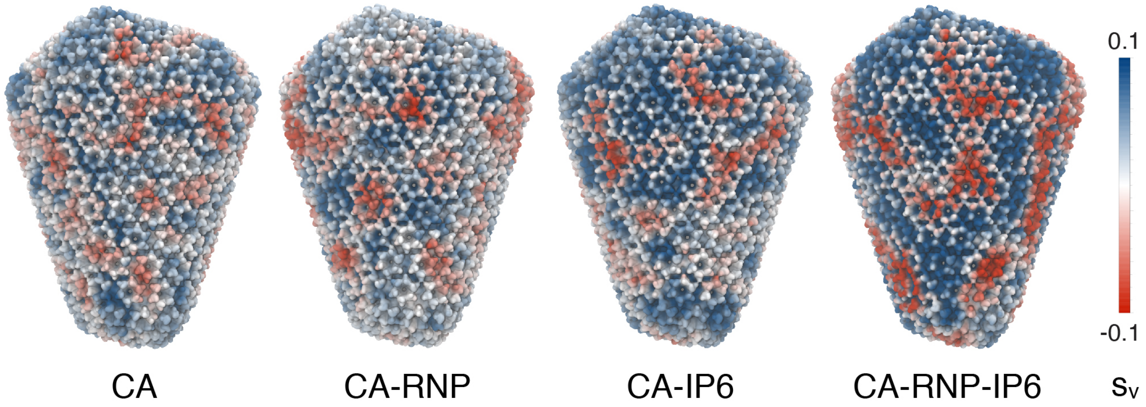


**Figure S2. Strain analysis of a HIV-1 core with a non-canonical morphology.** The volumetric strain was computed for an HIV-1 core containing liquid water, a model RNP complex, $IP_6$, and a RNP complex with $IP_6$. Red and blue colors correspond to compressive and expansive strain, respectively. Strain increased on the capsid upon the addition of RNP and $IP_6$, with CA < CA-RNP < CA-$IP_6$ < CA-RNP-$IP_6$.

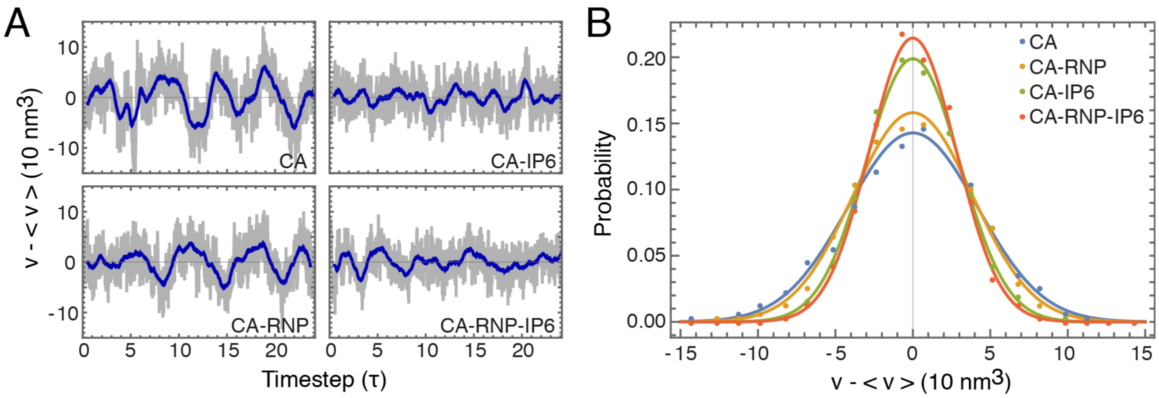


**Figure S3. Fluctuation analysis of a HIV-1 core with a non-canonical morphology.** **(A)** Mean-free volume fluctuations in the HIV-1 cores corresponding to the capsid morphology in fig. S1. The average core volume was $\langle V \rangle = 1.53 \times 10^5$ nm$^3$. The subsampled timestep, τ, is 4 ns. Gray colors denote the instantaneous volume, whereas the blue line denotes a moving average within a one-timestep window. **(B)** Probability distributions for the volume fluctuation amplitude across each system. Closed circles correspond to measured data, while the solid line indicates a Gaussian fit. The presence of the RNP and $IP_6$ increased capsid rigidity.

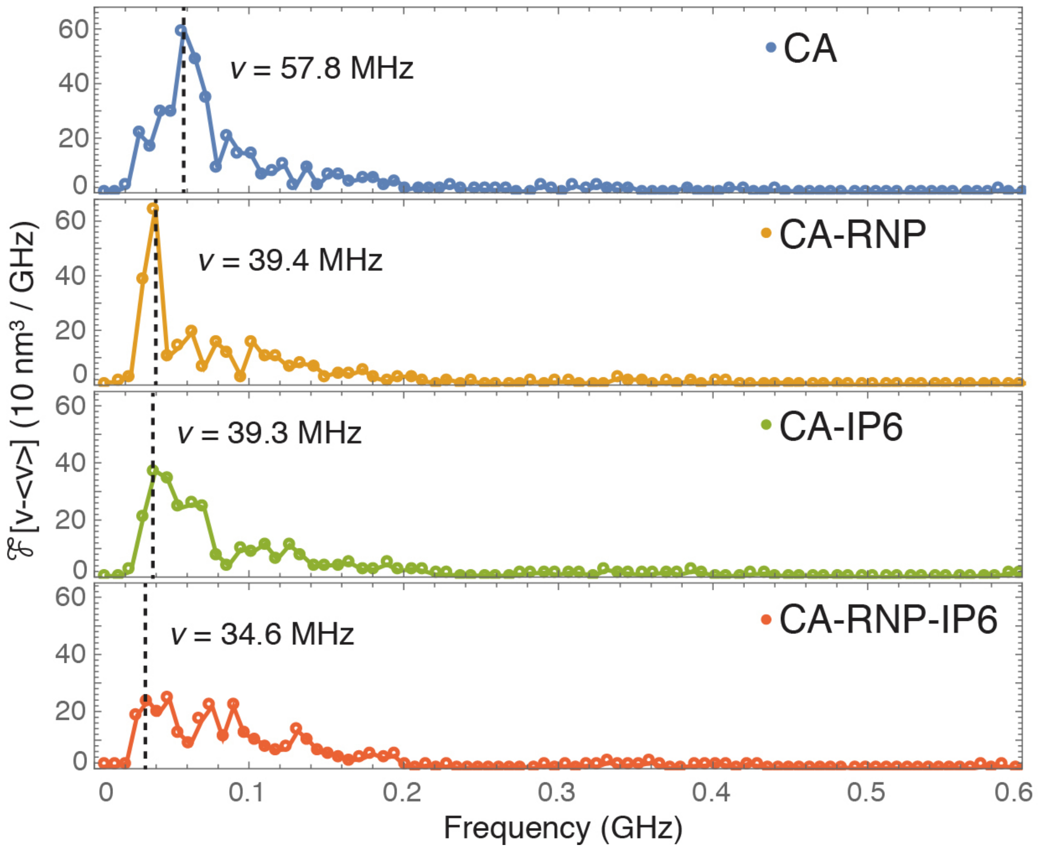


**Figure S4. Fourier analysis of the volume fluctuations.** The Fourier transform was computed for the volume fluctuations ($\mathcal{F}[V - \langle V \rangle]$) for the capsid containing liquid water, RNP, $IP_6$ and $IP_6$ with RNP. The dominant fluctuation mode shifts towards lower frequencies in the presence of the RNP (CA: $\upsilon = 57.8$ MHz; CA-RNP: $\upsilon = 39.4$ MHz), whereas $IP_6$ binding causes a broadened spectrum with larger intensities at higher frequency modes.

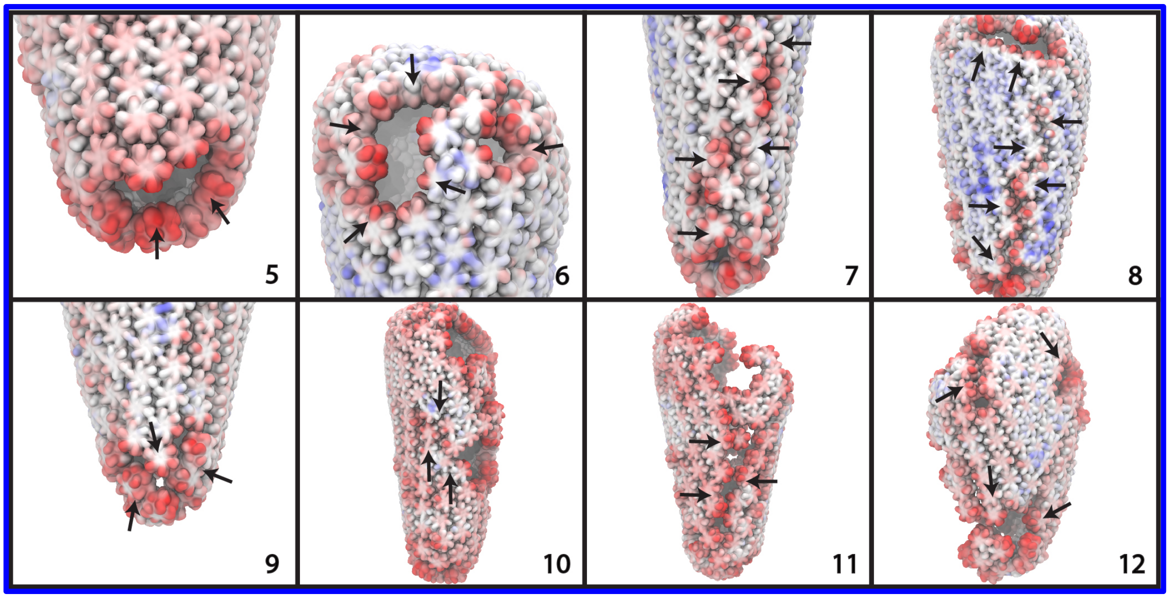


**Figure S5. Defect types experimentally observed in ruptured core lattices.** Expanded view of the ruptures, holes, and large cracks found in cores undergoing reverse transcription at advances stages of capsid disassembly. Lattice separations found in these structures are highlighted with black arrows. Capsids are numbered as in Fig. 5.

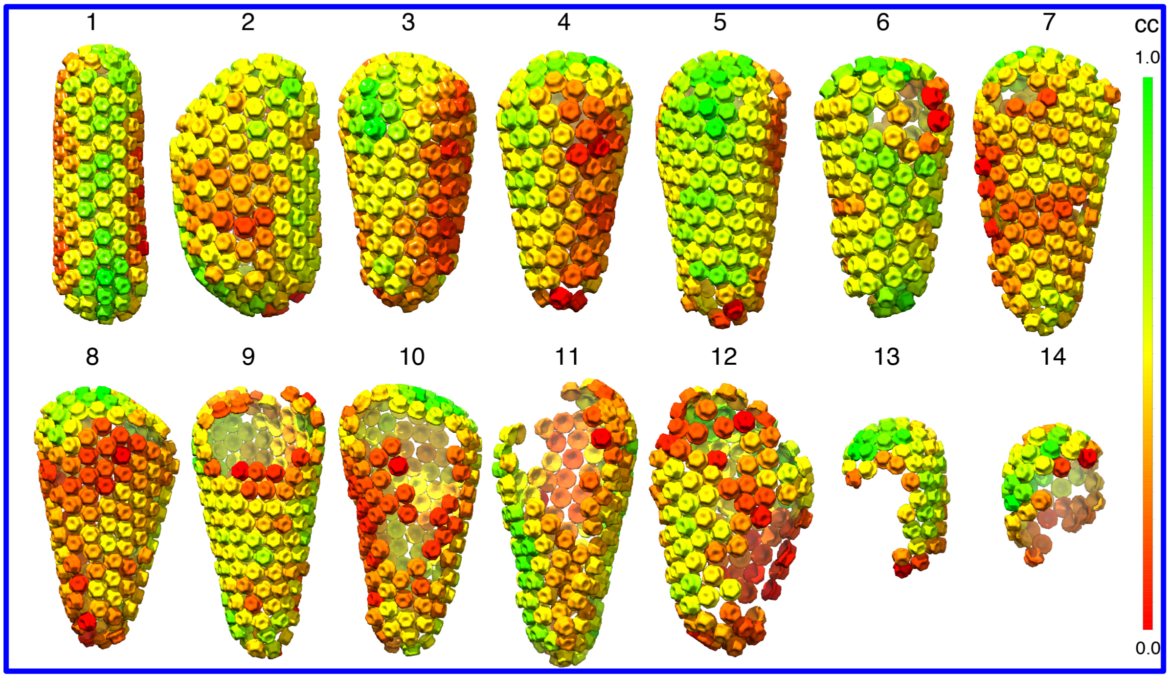


**Figure S6. Cross-correlation of the CA lattice with the cryo-tomograms.** The uncertainty in CA placement is determined by computing the cross-correlation (cc) normalized across each core between a cryo-EM map of the CA hexamer (EMD-3465) downsampled to 25 Å resolution and the local density in the collected subtomograms. The uncertainty values do not correlate with the lattice separation order parameter ($\chi$).

| System | Number of CA domains | Containing | Time simulated (ns) |
|---|---|---|---|
| Capsid 1 | 1308 | Water | 150.8 |
| Capsid 1 | 1308 | $IP_6$ | 150.8 |
| Capsid 1 | 1308 | RNP | 150.8 |
| Capsid 1 | 1308 | $IP_6$, RNP | 150.8 |
| Capsid 2 | 1242 | Water | 114.5 |
| Capsid 3 | 1200 | Water | 101.8 |
| Capsid 4 | 1146 | Water | 105.8 |
| Capsid 5 | 1116 | Water | 105.5 |
| Capsid 5 | 1116 | $IP_6$ | 131.1 |
| Capsid 6 | 1260 | Water | 126.6 |
| Capsid 6 | 1260 | $IP_6$ | 126.6 |
| Capsid 6 | 1260 | RNP | 126.6 |
| Capsid 6 | 1260 | $IP_6$, RNP | 126.6 |

**Table S1:** All-atom MD simulations of the HIV cores derived from cryo-ET in differing configurations containing water, $IP_6$, the RNP complex. The morphologies used for each capsid are numbered 1–6 corresponding to the structures from left to right in Fig. 1 A. CA hexamers and pentamers used in the initial configuration were modeled after the cryo-ET structure derived from intact virions (PDB ID: 5MCX, 5MCY).
